# Comparative transcriptomics reveals the molecular toolkit used by an algivorous protist for cell wall perforation

**DOI:** 10.1101/2021.12.30.474559

**Authors:** Jennifer V. Gerbracht, Tommy Harding, Alastair G. B. Simpson, Andrew J. Roger, Sebastian Hess

## Abstract

Microbial eukaryotes display a stunning diversity of feeding strategies, ranging from generalist predators to highly specialised parasites. The unicellular “protoplast feeders” represent a fascinating mechanistic intermediate, as they penetrate other eukaryotic cells (algae, fungi) like some parasites, but then devour their cell contents by phagocytosis. Besides prey recognition and attachment, this complex behaviour involves the local, pre-phagocytotic dissolution of the prey cell wall, which results in well-defined perforations of species-specific size and structure. Yet, the molecular processes that enable protoplast feeders to overcome cell walls of diverse biochemical composition remain unknown. We used the flagellate *Orciraptor agilis* (Viridiraptoridae, Rhizaria) as a model protoplast feeder, and applied differential gene expression analysis to examine its penetration of green algal cell walls. Besides distinct expression changes that reflect major cellular processes (e.g. locomotion, cell division), we found lytic carbohydrate-active enzymes that are highly expressed and upregulated during the attack on the alga. A putative endocellulase (family GH5_5) with a secretion signal is most prominent, and a potential key factor for cell wall dissolution. Other candidate enzymes (e.g. lytic polysaccharide monooxygenases) belong to families that are largely uncharacterised, emphasising the potential of non-fungal micro-eukaryotes for enzyme exploration. Unexpectedly, we discovered various chitin-related factors that point to an unknown chitin metabolism in *Orciraptor*, potentially also involved in the feeding process. Our findings provide first molecular insights into an important microbial feeding behaviour, and new directions for cell biology research on non-model eukaryotes.

## Results and discussion

### Food acquisition in Orciraptor agilis captured by comparative transcriptomics

*Orciraptor agilis* is a flagellate of the family Viridiraptoridae (Rhizaria) that can feed on the cell contents of dead algal cells (necrophagy)^1^. After attachment to its prey, filaments of *Mougeotia* sp. (Zygnematophyceae, Streptophyta), it perforates the algal cell wall and phagocytoses the nutrient-rich chloroplast (Figure 1A). The cell wall dissolution is confined to a narrow elliptical zone, which results in removal of a lid-like cell wall disc (Figure 1B). This perforation pattern appears to be defined by a transient F-actin-rich domain, the lysopodium^2^ (Figure 1C). As revealed by scanning electron microscopy, *Orciraptor* degrades both main structural components of the plant-like algal cell wall, (i) crystalline cellulose microfibrils and (ii) gel-like pectic substances (Figure 1D). No mechanical systems for cell wall perforation have been observed at an ultrastructural level. Instead, the close contact of *Orciraptor*’s plasma membrane to the zone of cell wall erosion indicates contact digestion^2^ (Figure 1C).

**Figure 1:**
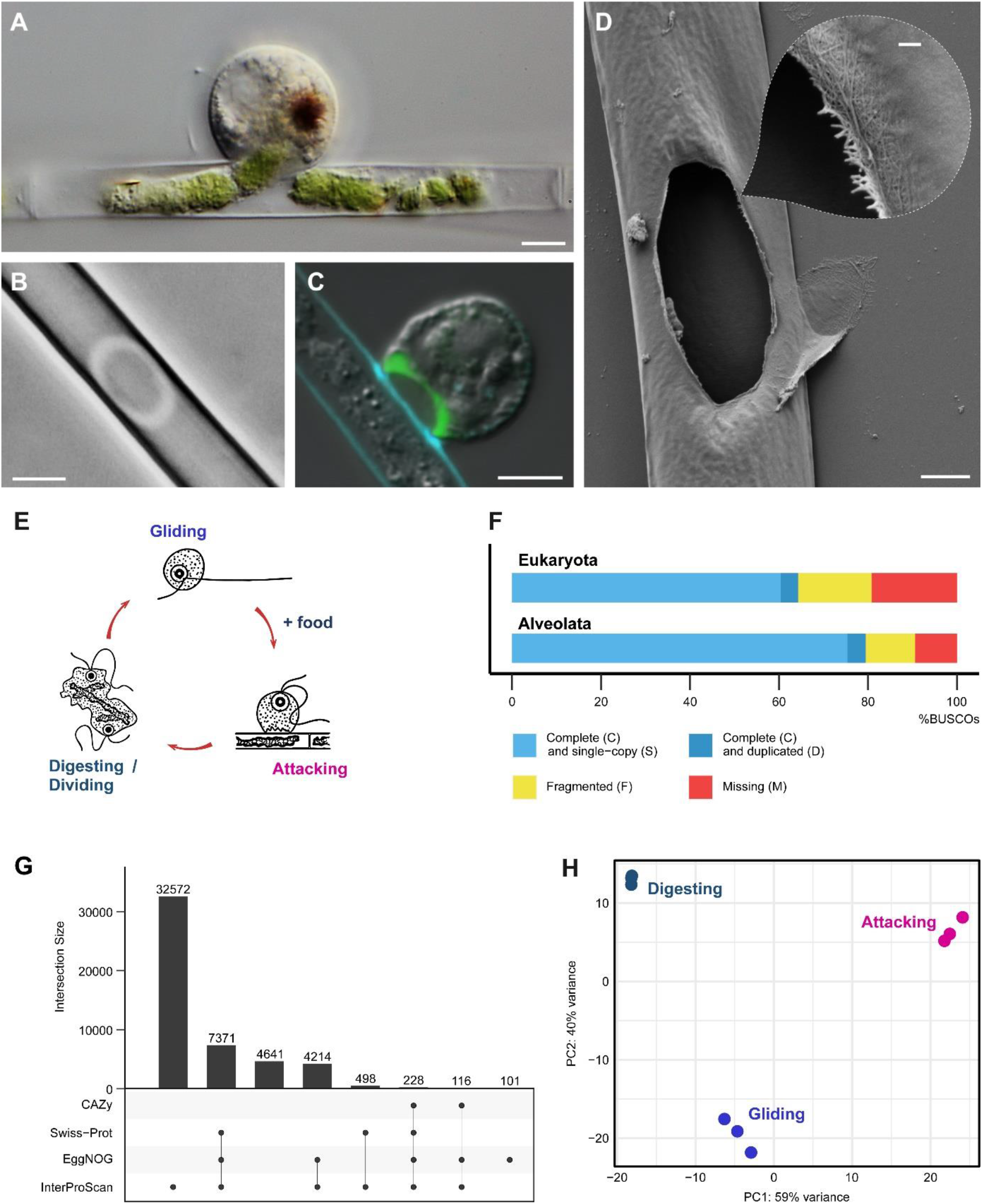
Feeding, life history and *de novo* transcriptome assembly of *Orciraptor agilis*. **A:** *Orciraptor agilis* extracting the chloroplast of *Mougeotia* sp. after perforating the algal cell wall (differential interference contrast). Scale bar 5 μm **B:** Annular dissolution of the algal cell wall resulting from an attempted attack (phase contrast). Scale bar 5 mm. **C:** Distribution of F-actin (green: fluorescent phalloidin) reveals the lysopodium in *Orciraptor* formed during attack on *Mougeotia* (overlay of differential interference contrast and fluorescence channels). The increased blue fluorescence (Calcofluor White) at the contact sites indicates lysis of the algal cell wall. Scale bar 5 μm. **D**: Scanning electron micrograph of a perforation by *Orciraptor* reveals the degradation of both main structural components of *Mougeotia’s* cell wall, gel-like biopolymers (potentially pectins, smooth surface) and cellulose microfibrils. Scale bars 2 μm and 200 nm (inset). **E**: Life history stages of *Orciraptor agilis* from which the samples were generated. **F**: BUSCO (benchmarked universal single-copy orthologs) assessment of the assembled transcriptome. The analysis was performed with the „Eukaryota” dataset and the „Alveolata” dataset (sister group of Rhizaria). **G**: Upset plot showing the number and overlap of ORFs annotated by the indicated annotation tools and databases. Only intersection sizes > 100 are shown. **H**: Principal-component analysis (PCA) based on the expression level of all transcripts for each replicate included in the experiment.

To gain insight into the molecular mechanisms underlying this feeding process, we compared the transcriptomes of *Orciraptor* cultures in three well-defined life history stages (= conditions, Figure 1E): 1) motile, gliding flagellates searching for algal cells (“gliding”), 2) cells during cell wall perforation about 45 min after contact with algal cells (“attacking”), and 3) a culture with excess algal material, which was enriched in digesting and dividing cells (“digesting-dividing”). *Orciraptor* is an excellent laboratory model, as it can be synchronised by starvation and attacks within a few minutes after addition of algal cells. Both its ability to grow under bacteria-free conditions and its preference for dead algae let us observe *Orciraptor*’s gene expression changes very clearly (no bacterial transcripts, no adaption by the algal food). Since there is no high-throughput genomic or transcriptomic data available for the Viridiraptoridae, we generated a transcriptome assembly *de novo* using the data from all conditions (nine samples, plus a sample of *Mougeotia* sp. to identify algal reads). This assembly captures the most complete picture of the transcriptomic landscape and was later used for read mapping. The transcriptome was determined to be 64.3% and 80.5% complete as assessed with BUSCO using Eukaryota and Alveolata datasets, respectively (Figure 1F). These values likely underestimate the true completeness substantially, given the relatively isolated phylogenetic position of *Orciraptor*. Using four different tools for functional annotation (dbCan, DIAMOND/Swiss-Prot, eggNOG-mapper, InterProScan), 90.7% of the 49,848 predicted open reading frames (ORFs) could be annotated (Figure 1G and Table S1). A principal component analysis of all replicates showed tight and highly distinct clusters for each condition (Figure 1H).

### Global expression changes reflect Orciraptor’s life history

To explore cellular processes affected by transcriptional changes between the life history stages of *Orciraptor*, we performed a differential expression analysis for each pair of two conditions. Differentially expressed transcripts (|log2 fold change| > 1, adjusted p-value < 0.001) in either of these comparisons were hierarchically clustered based on their relative expression changes in all conditions (Figure 2A). Applying an 80% maximum-height cut-off criterion to the clustering dendrogram yielded five clusters containing transcripts with similar expression patterns (Figure 2B). For each cluster, significantly enriched GO terms were determined (Figure S1), thereby identifying cellular processes associated with marked expression changes during *Orciraptor*’s life history. In addition, we specifically investigated expression changes in transcripts for cytoskeletal proteins (e.g. actin- and microtubule-related factors, motor proteins), as viridiraptorids shift from a rigid flagellate to an amoeboid stage during feeding.

**Figure 2:**
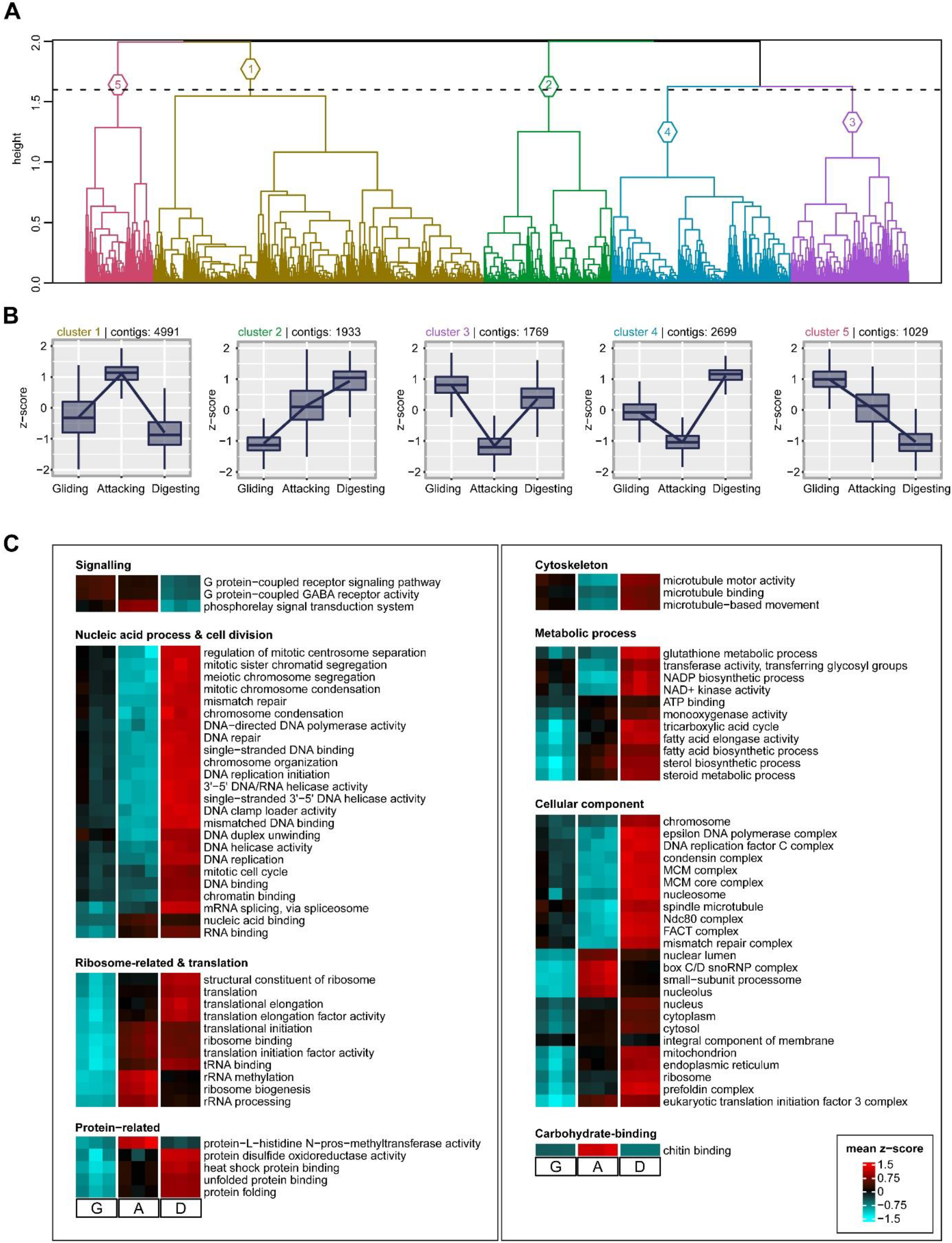
Clustering of differentially expressed transcripts and expression changes throughout *Orciraptor’s* life history. **A**: Hierarchical clustering of transcripts that were differentially expressed (|log2 fold change| > 1, adjusted p-value < 0.001) in at least one pairwise comparison. Variance stabilising transformed counts were used to perform hierarchical cluster analysis using Pearson correlation as distance method and complete linkage. The resulting dendrogram was cut at 80% of the maximum height, yielding five clusters of transcripts with similar expression patterns which are shown in (**B**). **C**: Heatmap of differentially expressed transcripts belonging to the GO terms that were significantly enriched in any of the five clusters. G: Gliding, A: Attacking, D: Digesting. Each square represents one biological replicate.

Gliding cells show relatively low expression levels in most of the listed terms and some categories appear to be specifically down-regulated (Figure 2C). This includes terms related to protein production such as ribosome biogenesis, translation, and RNA-related processes (splicing, binding), as well as some important metabolic processes (TCA cycle, fatty acid biosynthesis, sterol biosynthesis). This aligns well with the fact that gliding cells move around, but do not eat; they are probably in an “energy-saving mode” until food is encountered. Interestingly, we found a few kinesin homologues that are specifically upregulated in the “gliding” condition and are most closely related to flagellum-associated kinesins, e.g. a KIF17/OSM-3 homologue, a member of the kinesin-2 family of plus-end directed microtubule-based motor proteins (Figure S2, asterisk). This family is well studied in connection with intraflagellar transport (IFT) in *Chlamydomonas*^3^. *Orciraptor* performs a gliding motility that apparently relies on a traction system located in the adhering posterior flagellum^1^. This form of motility is widespread among heterotrophic flagellates of various phylogenetic affinities, but poorly understood. The locomotion in *Orciraptor* might be driven by an anterograde membrane motion along this flagellum, which in IFT is based on kinesins.

In the “attacking” condition, gene categories associated with ribosomal RNA and ribosome biosynthesis, protein production and some energy- and lipid-related metabolic processes show enhanced expression, indicating a marked switch in cellular activity upon contact with the algal cells. The pronounced expression of genes linked to ribosomal RNA processing and ribosome assembly during attack suggests that the protein production machinery becomes restored to full capacity in preparation for upcoming cellular processes such as phagocytosis, digestion, synthesis of biomass, and multiplication. Furthermore, the “attacking” condition is characterised by a high and specific expression of various myosins (Figure S2), some of which might be involved in the formation and maintenance of pseudopodial structures. Upon contact with algal cells, *Orciraptor* switches from a motile microtubule-dominated, flagellate to an F-actin-dominated, amoeboid cell. In this amoeboid stage, *Orciraptor* develops the’lysopodium’ as a cytoskeletal template for cell wall perforation^2^ and later uses pseudopodia to extract algal cell contents^1^. Interestingly, transcripts corresponding to the term “chitin binding” are highly upregulated during attack (Figure 2C), although the green algal food is unlikely to contain any chitinous substances (see below for details).

In the “digesting-dividing” condition, transcripts associated with translation and protein production remain highly expressed – similar to the “attacking” condition, yet with a slightly different pattern. However, there were also some profound and specific expression changes in the “digesting-dividing” condition. Transcripts related to signalling show the lowest expression levels of all studied life history stages, while energy conversion, lipid biosynthesis and glutathione-related processes were at maximum expression (Figure 2C). This may reflect the conversion of algal chloroplast material into viridiraptorid biomass that happens during the digestive phase. Glutathione-related processes, in particular, might be involved in the detoxification of ingested algal food, as chlorophylls and their breakdown products are known to produce reactive oxygen species when exposed to light^4^. A very pronounced upregulation in our global analysis is observed in transcripts related to DNA dynamics and cell division. Detailed examination of the expression of cytoskeletal components revealed a large fraction (two-thirds) of kinesins that are specifically upregulated during the “digesting-dividing” stage, together with regulators of chromosome condensation (RCC1), mitotic spindle-associated factors (ASPM) and centrin (Figure S2). This matches the observation that viridiraptorids only undergo mitosis and divide after food uptake^1^, while “gliding cells” seem to be arrested in a pre-division stage, as evidenced by the presence of probasal bodies for flagellar duplication^5^. In viridiraptorids, both mitosis and cytokinesis rely heavily on microtubular structures, the spindle apparatus and a cortical system of overlapping cytoplasm microtubules^2^, which may explain the marked expression changes of microtubule-related factors. All in all, our transcriptomic data clearly reflect the main cellular processes observed during the three studied life history stages of *Orciraptor*.

### Lytic CAZymes are highly upregulated during attack

The conjugating green alga *Mougeotia* sp. possesses a cell wall with certain similarities to those of the closely related land plants, including structural components such as crystalline cellulose microfibrils and gel-like pectins^6,7^. We suspected that *Orciraptor* utilises carbohydrate-active enzymes (CAZymes) to degrade these polymers (e.g. cellulases and pectinases) and analysed the expression of annotated lytic CAZymes such as glycoside hydrolases (GH) and polysaccharide lyases (PL). *Orciraptor* expressed a great diversity of GHs as well as some PLs listed in Figure 3A according to their expression level in the “attacking” condition. Indeed, there are several GHs with putative cellulase activity (GH5_5, GH5, GH6, GH44) as well as GHs or PLs that might degrade pectin or pectate (GH28 mainly contains polygalacturonases^8^; PL9_2 is a subfamily of PL9, whose members break up homogalacturonan by a β-elimination mechanism^9^). Some of these candidates were clearly upregulated in the “attacking” condition, especially the members of families GH5_5 and PL9_2 (Figure 3A).

**Figure 3:**
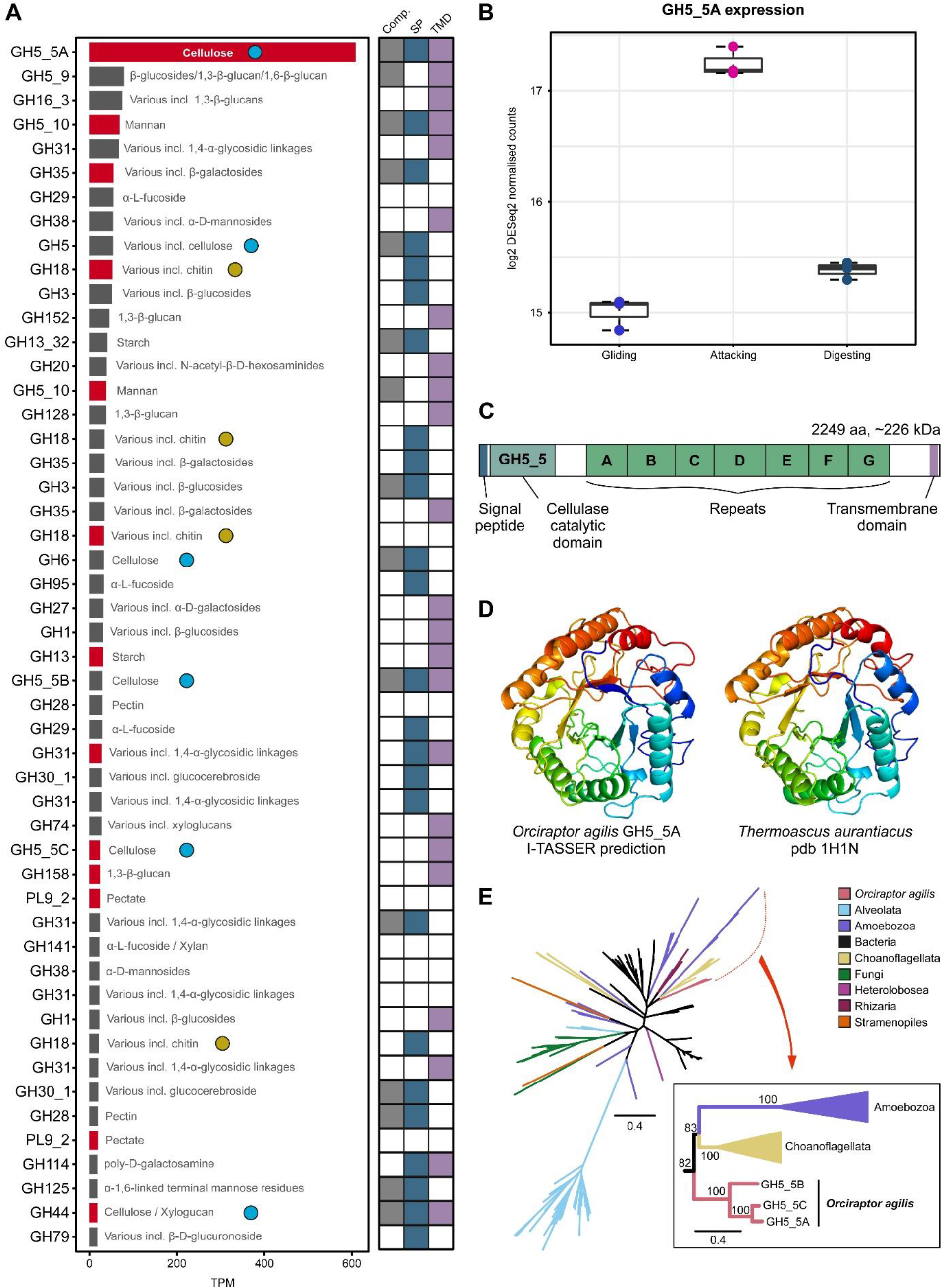
Glycoside hydrolases and polysaccharide lyases of *Orciraptor agilis*, with details on a highly expressed putative endocellulase. **A**: Top 50 most highly expressed CAZymes of the glycoside hydrolase (GH) and polysaccharide lyases (PL) families in the “attacking” condition. Expression levels are shown as transcripts per million (TPM). A red bar indicates upregulation (log2 fold change > 1, adjusted p-value< 0.001) in the attacking versus “gliding” condition. The main substrates of the respective CAZyme families are listed. The coloured boxes indicate whether the contig is complete (“Comp.”, grey), and has a signal peptide (SP, blue) or transmembrane domains (TMD, purple). Contigs annotated as CAZymes with putative endoglucanase function (EC 3.2.1.4) are marked with a light blue dot. Contigs annotated as CAZymes that target chitin are marked with a yellow dot. **B**: Expression levels as normalised counts of the most highly expressed GH5_5 contig (GH5_5A). Each dot represents one biological replicate. **C**: Schematic depiction of the GH5_5A functional domains. **D**: *In silico* structure prediction of the GH5_5 domain from *Orciraptor agilis* GH5_5A shown next to an endoglucanase from *Thermoascus aurantiacus* **E**: Radial phylogenetic tree of GH5_5 family proteins from bacteria and eukaryotes. Highlighted are the three GH5_5 sequences from *Orciraptor agilis*. Ultrafast bootstrap values are shown as branch support.

The most highly expressed CAZyme by far, here termed GH5_5A, showed a marked and specific upregulation in the “attacking” condition, with a log2 fold change of 2.2 (adjusted p-value 3.1 × 10^−136^) compared to the “gliding” condition (Figure 3B). The contig was split into two in the original assembly, most likely due to intronic sequences present (Figure S3A). It was extended and completed by using an alternative assembly strategy (see Methods for details).

Characterised family GH5_5 members from other organisms are typical endocellulases, i.e. they perform internal cleavage of β(1->4) glucosidic linkages^10^. This activity is also required for the degradation of cellulose microfibrils in plant-like cell walls, making the highly expressed and regulated GH5_5A of *Orciraptor* a strong candidate for an important wall-degrading role. The complete ORF of GH5_5A is 2249 amino acids long, which corresponds to a large protein of approximately 226 kDa with an N-terminal signal peptide and a C-terminal transmembrane domain (TMD, Figure 3C). The protein might be secreted and remain tethered to a membrane. The GH5_5 domain is followed by a series of seven related sequence motifs, some of which are weakly assigned to the cellulose-binding domain CBM2 (with a non-significant E-value). Future wet-lab studies will be required to elucidate whether these repeats possess the ability to bind cellulose. Based on these features, it is possible that the enzyme is anchored on the external side of the plasma membrane and aids in contact digestion when *Orciraptor* is attached to its prey.

To gain more insight into the catalytic function of GH5_5A, we predicted *in silico* the structure of the GH5_5 module using the I-TASSER service (Iterative Threading ASSEmbly Refinement^11-13^). The predicted tertiary structure showed a high similarity to the experimentally determined atomic resolution structure of the “major endoglucanase” from the fungus *Thermoascus aurantiacus* (Figure 3D)^14^. Furthermore, a multiple sequence alignment of the GH5_5 modules from *Orciraptor agilis* and several endocellulases of the same family revealed conservation of residues that are part of the active site of a functionally and structurally characterised GH5_5 endoglucanase (Figure S3B)^15,16^.

We found two other transcripts that encode GH5_5 modules in *Orciraptor*, GH5_5B and GH5_5C. Both had much lower expression levels, and only GH5_5C was upregulated in the “attacking” condition.

To elucidate the relationships of the three endocellulases of *Orciraptor*, we performed phylogenetic analyses with prokaryotic and eukaryotic GH5_5 homologues. The maximum likelihood tree in Figure 3E shows that GH5_5 sequences from diverse eukaryotic supergroups do not cluster together but are intermingled with prokaryotic sequences. Fungal and dinoflagellate (Alveolata) cellulases, however, form two distinct clades indicating significant in-group diversification (Figure 3E), which might relate to their cell wall biology (e.g. cellulosic thecal plates in dinoflagellates^17^). The three GH5_5 domains from *Orciraptor* clustered together as well, with full bootstrap support, suggesting that they are paralogs stemming from a common ancestral gene. Their closest relatives in the tree were sequences from choanoflagellates, but due to lacking statistical support and poor taxon sampling of eukaryotes in general^18^, the origin of *Orciraptor*’s GH5_5 cellulases remains unresolved. Future efforts in the genomic exploration of microbial eukaryotes promise to fill these gaps and to provide further evolutionary insights.

### Are chitin-related factors involved in contacting algal surfaces?

The analysis of *Orciraptor*’s glycoside hydrolases also revealed several putative chitinases from the GH18 family, some of which were upregulated in the “attacking” condition (Figure 3A). This was surprising, as conjugating green algae such as *Mougeotia* sp. do not produce detectable chitin, nor do they possess chitin synthases in their genomes^19^. Another possibility is that chitin plays a role in the physiology of *Orciraptor*, especially during feeding. Noting the marked expression changes of “chitin binding” factors in our global analysis (Figure 2C), we examined carbohydrate-binding modules (CBMs) and their expressional changes. *Orciraptor* expressed proteins with various CBMs as listed in Figure 4A according to their expression level in the “attacking” condition. The most highly expressed CBMs belong to family CBM13, which contains members with diverse binding functions (galactose and mannose residues, xylan, GalNAc and others), so that substrate specificity cannot be predicted^20^. Interestingly, there were several transcripts with CBM50 (LysM) and/or CBM18 modules, both of which are known to bind chitin (and peptidoglycan)^21,22^. Several of these factors were upregulated during attack as well. The most highly expressed and upregulated chitin-binding transcript encodes five CBM50 modules and a single CBM18 module (Figure 4B). It also has a signal peptide and a C-terminal transmembrane domain, and, hence, could be tethered to a membrane, similar to known LysM-containing chitin receptors in plants^23^. These findings are a good starting point for future research, as LysM domains are important factors in plant-pathogen interactions and rhizobial symbiosis^24^, but largely unexplored in free-living protists. In addition to chitinases and putative chitin-binders, we found other chitin-related factors that were relatively highly expressed, such as potential lytic polysaccharide monooxygenases (LPMO) of family AA11 and a chitin synthase (Figure 4C). The array of chitin-synthesising, -binding and -degrading factors in *Orciraptor* points to a significant role of chitin (or related substances). Indeed, early analytical results indicate that *Orciraptor* produces and secretes chitinous material during attack on algal cells (work in progress), which might explain the differential expression of chitin-related factors. The properties and roles of such biopolymers in protists are largely unknown and deserve future study.

**Figure 4:**
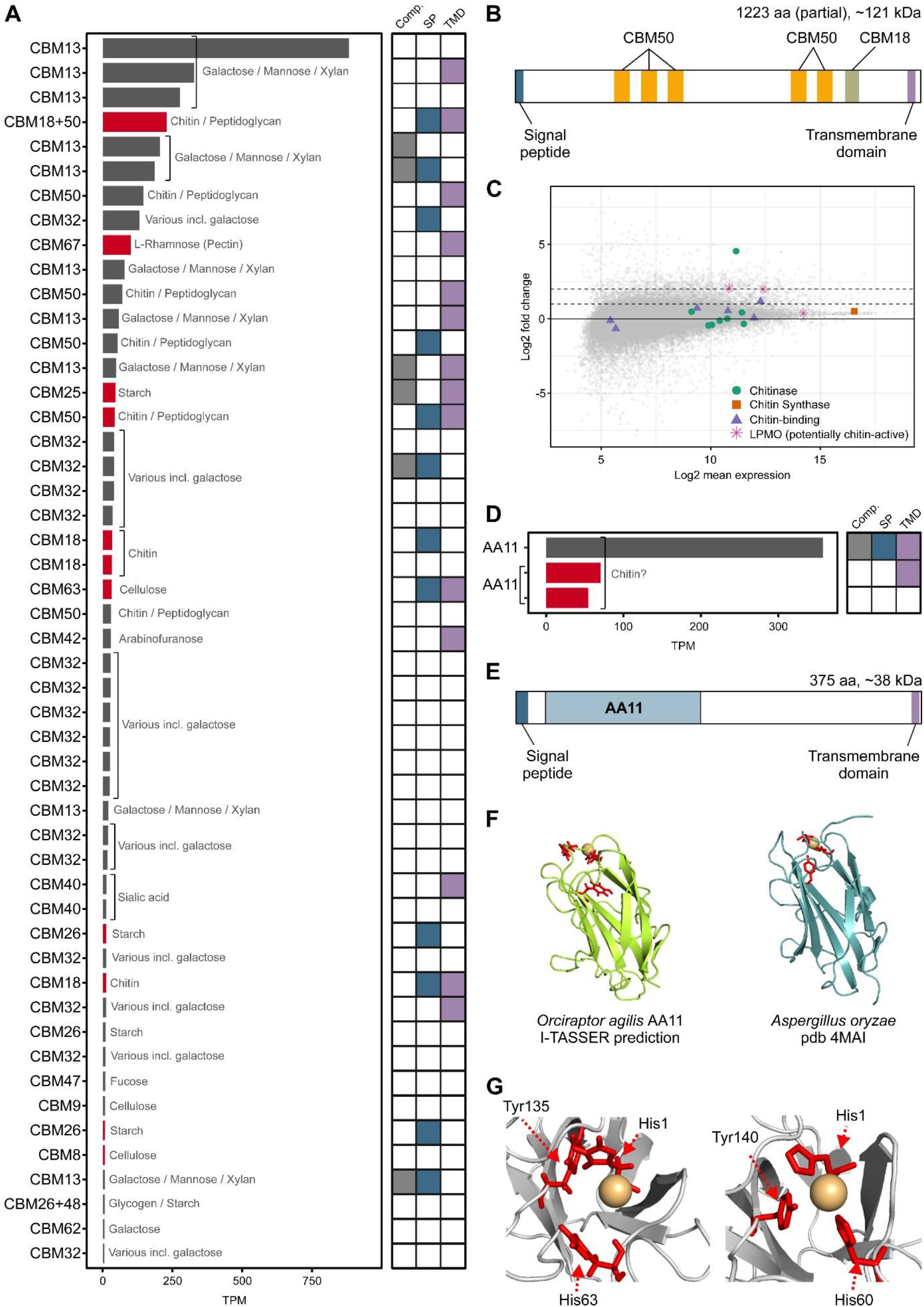
Carbohydrate-binding modules, lytic polysaccharide monooxygenases and other chitin-related factors expressed in *Orciraptor agilis*. **A**: Top 50 most highly expressed contigs annotated with carbohydrate-binding modules (CBMs). Expression levels are shown as transcripts per million (TPM). A red bar indicates upregulation (log2 fold change > 1, adjusted p-value < 0.001) in the “attacking” versus “gliding” condition. The targeted carbohydrates of the respective CBMs are listed. The coloured boxes indicate whether the contig is complete (“Comp.”, grey), and has a signal peptide (SP, blue) or transmembrane domains (TMD, purple). **B**: Schematic depiction of the most highly expressed chitin-related contig. **C**: Volcano plot depicting the expression levels of contigs annotated with a chitin-related function. **D**: Expression levels of contigs annotated as AA11 shown as transcripts per million (TPM). A red bar indicates upregulation (log2 fold change > 1, adjusted p-value < 0.001) in the “attacking” versus “gliding” condition. The coloured boxes indicate whether the contig is complete (“Comp.”, grey), and has a signal peptide (SP, blue) or transmembrane domains (TMD, purple). **E**: Schematic depiction of the most highly expressed AA11 from *Orciraptor agilis*. **F**: *In silico* structure prediction of the functional domain of the AA11-type LPMO from *Orciraptor agilis* next to the AA11 LPMO from *Aspergillus oryzae* (residues 1 – 151). The copper cofactor (orange) and three binding residues (red) in the *Orciraptor* structure were predicted by COACH and COFACTOR. **G**: Details of the ligand binding sites of the structures shown in (**F**).

Three contigs encode putative LPMOs of family AA11 (Figure 4D). LPMOs are very promising enzymes for biotechnology as they act on recalcitrant substrates such as crystalline cellulose or chitin, and greatly enhance biomass degradation in synergy with other CAZymes^25,26^. However, only a single characterised enzyme is known for family AA11, the copper-dependent LPMO from *Aspergillus oryzae* that degrades chitin^27^. The most highly expressed putative LPMO from *Orciraptor* encodes a 375 amino acid long ORF, again with an N-terminal signal peptide and a C-terminal transmembrane domain (Figure 4E). The sequence similarity between *Orciraptor*’s LPMO module and the module of the characterised AA11 protein is relatively low (25.4% identity), but their relationship is supported by *in silico* structure prediction. The AA11 LPMO from *Aspergillus oryzae* was the closest hit as determined with I-TASSER, and clearly resembles the predicted structure from *Orciraptor* (Figure 4F). Furthermore, protein-ligand binding site predictions with COACH^28^ and COFACTOR^29-31^ predicted a divalent copper ion as ligand, which is typical for known LPMOs^32^. The ligand was bound by a trio of amino acid residues (His1, His63, Tyr135) that is very similar to that known to bind copper in the AA11 of *Aspergillus* (Figure 4G). We do not yet know the activity of the putative LPMO from *Orciraptor*, nor its role in *Orciraptor*’s biology, but our finding demonstrates that non-fungal microeukaryotes represent an almost untapped resource for new enzymes with potential biotechnological relevance.

## Conclusion

Comparative transcriptomics applied to synchronised cultures of the protoplast feeder *Orciraptor* have provided the first insights into the molecular factors underpinning the perforation of algal cell walls. The pronounced upregulation of GH5_5A during attack, identifies this highly expressed glycoside hydrolase as a potential key factor for the pre-phagocytotic dissolution of the cell wall. The molecular features of this putative endocellulase (signal peptide, TMD) suggest that the protein is secreted but remains tethered to a membrane, e.g. the extracellular side of the plasma membrane. Interestingly, several other candidate proteins that were upregulated during attack (e.g. chitin-binders, LPMOs) display similar features, pointing to a membrane-tethered toolkit of CAZymes, similar to that known from bacterial cellulosomes^33^. Our findings support the hypothesis that protoplast feeders overcome prey cell walls by contact digestion, and that the mechanistic aspects of their intricate feeding strategy differ fundamentally from those of “typical” predators and fungal parasites. Other experimental approaches are now required to reveal more of the molecular secrets of protoplast feeders for which the sequence data generated in this study will be an important resource.

## Acknowledgements

This work was funded by the German Research Foundation grant 283693520 (to S.H.) and German Research Foundation grant 417585753 (to S.H.). Funding for the work carried out in A.J.R.’s laboratory was provided by a Discovery grant 2017-06792 from the Natural Sciences and Engineering Research Council (NSERC) of Canada. Funding for work in A.G.B.S.’s laboratory was supported by NSERC Discovery Grant 298366-2014. We thank Ruth Bruker (University of Cologne) for assistance with scanning electron microscopy.

## Author contributions

Conceptualisation: S.H.;

Investigation: J.V.G., T.H., and S.H.;

Writing - Original Draft: J.V.G., T.H., and S.H.;

Writing - Review & Editing: All authors;

Visualisation: J.V.G., S.H.;

Funding acquisition: A.G.B.S., A.J.R., and S.H.

## STAR Methods

### Resource availability

#### Lead contact

Further information and requests for resources and reagents should be directed to and will be fulfilled by the lead contact, Sebastian Hess.

#### Materials availability

This study did not generate new unique reagents.

#### Data and code availability

- RNA-seq data have been deposited at ArrayExpress and are publicly available as of the date of publication. Accession numbers will be listed in the key resources table.
- Transcriptome assemblies have been deposited at ENA and are publicly available as of the date of publication. Accession numbers will be listed in the key resources table.
- Peptide sequences, gene expression tables and functional annotation data have been deposited at Zenodo and are publicly available as of the date of publication. DOIs will be listed in the key resources table.
- All original code has been deposited at github and is publicly available as of the date of publication. DOIs will be listed in the key resources table.

### Experimental model and subject details

#### Mougeotia sp

The filamentous green alga *Mougeotia* sp. (strain CCAC 3626) was grown in vented polystyrene cell culture flasks (Falcon® T25; Corning, New York, USA) with the culture medium Waris-H containing 1 % (v/v) bacterial standard medium (0.8 % peptone, 0.1 % glucose, 0.1 % meat extract, 0.1 % yeast extract in distilled water (w/v); for references see Hess and Melkonian ^1^) and artificial light (white LEDs, photon fluence rate 10–30 μmol m^−2^ s^−1^, 14:10 h light-dark cycle) at 16 °C. The strain CCAC 3626 was deposited in and is available from the Central Collection of Algal Cultures (CCAC) at the University of Duisburg-Essen (https://www.uni-due.de/biology/ccac/).

#### Orciraptor agilis

*Orciraptor agilis* (strain OrcA03) was cultivated in a diluted suspension of freeze-killed filaments of *Mougeotia* sp. (strain CCAC 3626) at 4-21 °C as described previously^1^. In short, about 25 ml of an algal culture (details below) was mixed with about 475 ml sterile, distilled water, distributed to polystyrene cell culture flasks (e.g. 25 ml in Falcon® T25; Corning, New York, USA), frozen and stored at -20 °C. These flasks were thawed and then inoculated with about 2 ml of a running *Orciraptor* culture. The strain OrcA03 is available from the laboratory of the corresponding author.

### Method details

#### Microscopy

Light microscopy was done with a ZEISS IM35 inverted microscope equipped with differential interference contrast and phase contrast optics, and an electronic flash (details see Hess and Melkonian ^1^). For the localization of F-actin and algal cell walls, attacking *Orciraptor* cells were aldehyde-fixed, washed and stained with a fluorescent phalloidin conjugate and Calcofluor White as described in Busch and Hess ^2^. For scanning electron microscopy of cell wall perforations, filaments of *Mougeotia* emptied by *Orciraptor* were collected from old cultures by sedimentation and placed on poly-L-lysine coated cover slips. After about one hour of sedimentation, the cover slips were passed through a graded ethanol series (30%-50%-96%-100%; 5 min each step), transferred to 100% hexamethyldisilazane (HMDS) and incubated for 15 min. After a final exchange of HMDS, the fluid was aspirated and the samples were air-dried in a fume hood. The dry samples were sputter coated with gold and imaged with a ZEISS Neon 40 scanning electron microscope (secondary electron detector, 2.5 kV acceleration voltage; ZEISS, Oberkochen, Germany).

#### RNA isolation of Mougeotia

Algal filaments were collected with a 40 μm strainer (Falcon® 40 μm Cell Strainer; Corning, New York, USA) from a well-grown culture, added to liquid nitrogen in a ceramic mortar, and ground to powder. Several milliliters of TRIzol Reagent (Thermo Fisher Scientific Inc., Waltham, Massachusetts, USA) were added and mixed with the ground algal material during thawing. The resulting TRIzol extract was transferred in a test tube, mixed for several minutes at room temperature, and subjected to RNA isolation according to the manufacturer’s instructions of the TRIzol Reagent. The isolated RNA was checked for integrity by agarose gel electrophoresis, quantified, and stored frozen in nuclease-free water.

#### Synchronization of Orciraptor cultures and RNA isolation

Nine large cultures of *Orciraptor agilis* were set up in vented T-175 cell culture flasks (Sarstedt, Nümbrecht, Germany) by adding about 30 ml of an *Orciraptor* suspension (gliding, aggressive cells from regular cultures) to about 250 ml of freeze-killed, algal material diluted in distilled water (details on dilution above), and incubated at 21 °C in the dark. One day after inoculation, three cultures with digesting and dividing *Orciraptor* cells (“digesting-dividing” condition) were processed for extraction of total RNA (details below). Two days after inoculation, the remaining cultures contained gliding, aggressive flagellates. Three of these cultures were directly processed for RNA extraction (“gliding” condition), while the other three cultures were spiked with 250 μl concentrated, freeze-killed *Mougeotia* filaments. These algae have been ultrasonicated before, to fragment filaments into shorter pieces (for faster sedimentation), and washed two times with distilled water (by centrifugation at 1000 *g* for 10 min and resuspension). After about 45 min, when almost all *Orciraptor* cells had started attack on the *Mougeotia* filaments (“attacking” condition), total RNA was extracted.

For extraction of total RNA, cultures of all conditions were processed the same way: After careful aspiration of most of the culture supernatant, the cells were quickly agitated and filtered onto a 3 μm polycarbonate membrane disc filter (Sterlitech, Auburn, Washington, USA) with a vacuum filtration device. The cell-bearing filter was then put upside down in a 60 mm Petri dish with 3 ml TRIzol Reagent (Thermo Fisher Scientific Inc., Waltham, Massachusetts, USA), and placed on a rocking table. After several minutes of mixing, the filter was removed from the Petri dish and the TRIzol extract stored frozen at -80 °C until further processing. From these samples, RNA was isolated according to the manufacturer’s instructions of the TRIzol Reagent, then treated with the TURBO™ DNase (Thermo Fisher Scientific Inc., Waltham, Massachusetts, USA), checked for integrity by agarose gel electrophoresis, quantified, and stored frozen in nuclease-free water.

#### RNA-seq and de novo transcriptome assemblies

The RNA samples of *Mougeotia* and *Orciraptor* were submitted to Génome Québec (Montréal, Québec, Canada) for strand-specific library preparation (including poly-A-enrichment) and RNA sequencing on a HiSeq2500 platform (PE125) and HiSeq4000 (PE100), respectively. For *Mougeotia*, about 45 million read pairs were obtained. K-mer based error correction was performed with Rcorrector^34^ (version 1.0.4). Quality and adapter trimming was performed with Trim Galore^35,36^ (version 0.6.6). The processed reads were assembled using rnasSPAdes^37^ (version 3.15.0) using a strand-specific option (--ss rf).

For *Orciraptor*, about 392 million read pairs (30-50 million read pairs per sample) were generated. Rcorrector^34^ (version 1.0.4) was used to perform k-mer based error correction. Adapter and low-quality bases were trimmed with Trim Galore^35,36^ (version 0.6.6). The processed reads were mapped to ribosomal sequences of the SILVA SSU r138.1 database for the groups “Orciraptor” and “Mougeotia”. This step was performed using bowtie2^38^ (version 2.4.2) with the parameters --very-sensitive and --score-min C,0,0 and only unmapped reads were kept. These reads were then mapped to the *Mougeotia* transcriptome with bowtie2^38^ (version 2.4.2) using default parameters. Reads that did not map to the algal transcriptome were used for the *de novo* assembly.

Assembly with Trinity: The filtered reads from all three life history conditions of *Orciraptor* were pooled for a strand-specific (--SS_lib_type RF) *de novo* assembly using Trinity^39^ (version 2.0.6). The contigs were blasted against the nt database to detect potential contaminants (task: megablast, version 2.10.1). Sequences resulting in hits with > 95% identity over a length of minimum 100 nt that matched to ribosomal, algal, bacterial, or viral sequences were removed from the assembly. ORFs were predicted with TransDecoder^40^ (version 2.1.0). To remove redundancy in the assembly all ORFs, as protein sequences, were first compared to each other using DIAMOND^41^ (version 2.0.11). Then, for each pair sharing >95% identity along >90% of the shortest ORF in the pair, the longest ORF was kept for further analyses.

Assembly with rnaSPAdes: The assembly was also performed with rnaSPAdes^37^ (version 3.14.1) an a strand-specific way (--ss rf). This transcriptome was filtered for a minimum contig size of 200 nt. To identify potential contaminants, the remaining contigs were compared to the nt database using blastn (task: megablast, version 2.10.1). Contigs resulting in hits with > 95% identity over a length of minimum 100 nt that corresponded to ribosomal, algal, bacterial, or viral sequences were removed.

#### Assembly statistics

Transcriptome assembly statistics were obtained with the scripts “TrinityStats.pl” and “contig_ExN50_statistic.pl” from the Trinity^39^ toolkit utilities. The presence of single-copy orthologs was determined with BUSCO^42^ (version 4.0.6) for the lineage datasets “eukaryota_odb10” and “alveolata_odb10”. ORF statistics were obtained using the custom bash script transdecoder_count.sh.

#### Functional annotation

The predicted ORF sequences of the Trinity assembly were compared to the the UniProtKB/Swiss-Prot database (Release 2021_01) using DIAMOND^41^ (version 2.0.6). Furthermore, an InterProScan analysis^43^ (version 5.52-86.0) with lookup of corresponding pathway and Gene Ontology annotation was conducted (databases: CDD-3.18, Coils-2.2.1, Gene3D-4.3.0, Hamap-2020_05, MobiDBLite-2.0, PANTHER-15.0, Pfam-33.1, PIRSF-3.10, PIRSR-2021_02, PRINTS-42.0, ProSitePatterns-2021_01, ProSiteProfiles-2021_01, SFLD-4, SMART-7.1, SUPERFAMILY-1.75, TIGRFAM-15.0). EggNOG-mapper^44,45^ (version 2.0.5-6) was used in DIAMOND and HMM mode for a functional annotation based on precomputed orthologous groups and phylogenies. The results from the DIAMOND-based annotation were kept and HMM hits were added for sequences that were not annotated using DIAMOND. Carbohydrate-active enzymes were annotated with dbcan2^46,47^ (stand-alone version 3.0) in HMM mode using the dbCAN-HMMdb-V9 database. Transmembrane domains and signal peptides were predicted for selected protein sequences with the Phobius webserver^48^.

#### Differential expression analysis

The processed and filtered reads were mapped to the coding sequences obtained from the Trinity assembly with bowtie2^38^ (version 2.4.2). Transcript abundance was quantified with salmon^49^ (version 1.4.0) in alignment-based mode. The read counts were parsed with tximport^50^ (version 1.18.0) to generate matrices containing counts and abundances (TPM). A pre-filtering step only keeping contigs with CPM > 1 in 2 or more samples was applied. Differential expression analysis was performed with DESeq2^51^ (version 1.30.0).

#### Expression profiles and gene ontology enrichment analysis

Transcripts that were differentially expressed (|log2 fold change| > 1, adjusted p-value < 0.001) in at least one pairwise comparison were clustered based on variance stabilising transformed counts. Hierarchical clustering was performed using Pearson correlation as distance method and complete linkage. The resulting dendrogram was cut at 80% of the maximum height, yielding five clusters of transcripts with similar expression patterns. Gene Ontology (GO) terms were retrieved by mapping the DIAMOND/Swiss-Prot annotation and merging them with the ones retrieved in the InterProScan analysis in Blast2GO^52^ (version 5.2.5). GO term enrichment analysis was performed with GOseq^53^ (version 1.42.0). The sequence lengths required for the analysis were computed with the script “fasta_seq_length.pl” from the Trinity^39^ toolkit utilities.

#### Protein structure prediction and structure-based functional annotation

The structural modelling of the respective protein domains was performed using the I-TASSER web server^54^. PyMOL (version 1.8.x) was used for the visualisation of protein structures.

#### Extension of the GH5_5A contig

In the Trinity assembly, the predicted ORF of the most highly expressed GH5_5A was incomplete and lacked a stop codon. There was another ORF present with 100% identity in the GH5_5 module which might have been separated during assembly because of intronic sequences (see Figure S3A for predicted splicing sites). In the rnaSPAdes assembly seven isoforms for the GH5_5A cellulase belonging to one gene cluster were identified. To represent all transcripts per gene cluster, superTranscripts were constructed using Lace^55^ (version 1.14.1), reads were aligned to the superTranscriptome with STAR^56^ (version 2.7.8a) and sequences were visualised in IGV^57^ (version 2.9.2). The alignment was used in StringTie^58^ (version 2.1.5) to assemble transcripts. Next, the sequences were extracted using gffread^59^ and ORF prediction was performed with TransDecoder^40^. Using this approach, a GH5_5A transcript was extracted that encoded a complete ORF with a length of 2249 amino acids. Read mapping and differential gene expression analysis was repeated with the Trinity assembly in which the GH5_5A contigs were replaced by the extended sequence. This quantification was used for Figures 3A and B.

#### Phylogenetic analysis of GH5_5 domains

The GH5_5 domain sequence of *Orciraptor* GH5_5A was used as a query to search for homologs in the non-redundant NCBI database, as well as the EukProt database^60^. A multiple sequence alignment was created with MAFFT^61^ (version 7.487) applying the L-INS-i method. The alignment was trimmed with trimAl^62^ (version 1.4.rev15) using the -automated1 setting. Identical sequences from the trimmed alignment were removed in Jalview^63^ (version 2.11.1.4) using the “Remove Redundancy” function with a threshold of 100. The substitution model with the best-fit was determined to be Q.pfam+R5 by the ModelFinder^64^ function of IQ-TREE^65^ (version 2.1.4-beta). A maximum likelihood tree was computed with IQ-TREE using this model and branch support values were calculated with UFboot^66^ with 1000 bootstrap replicates. The tree was visualised with FigTree (version 1.4.4).

## Supplemental information

**Figure S1:**
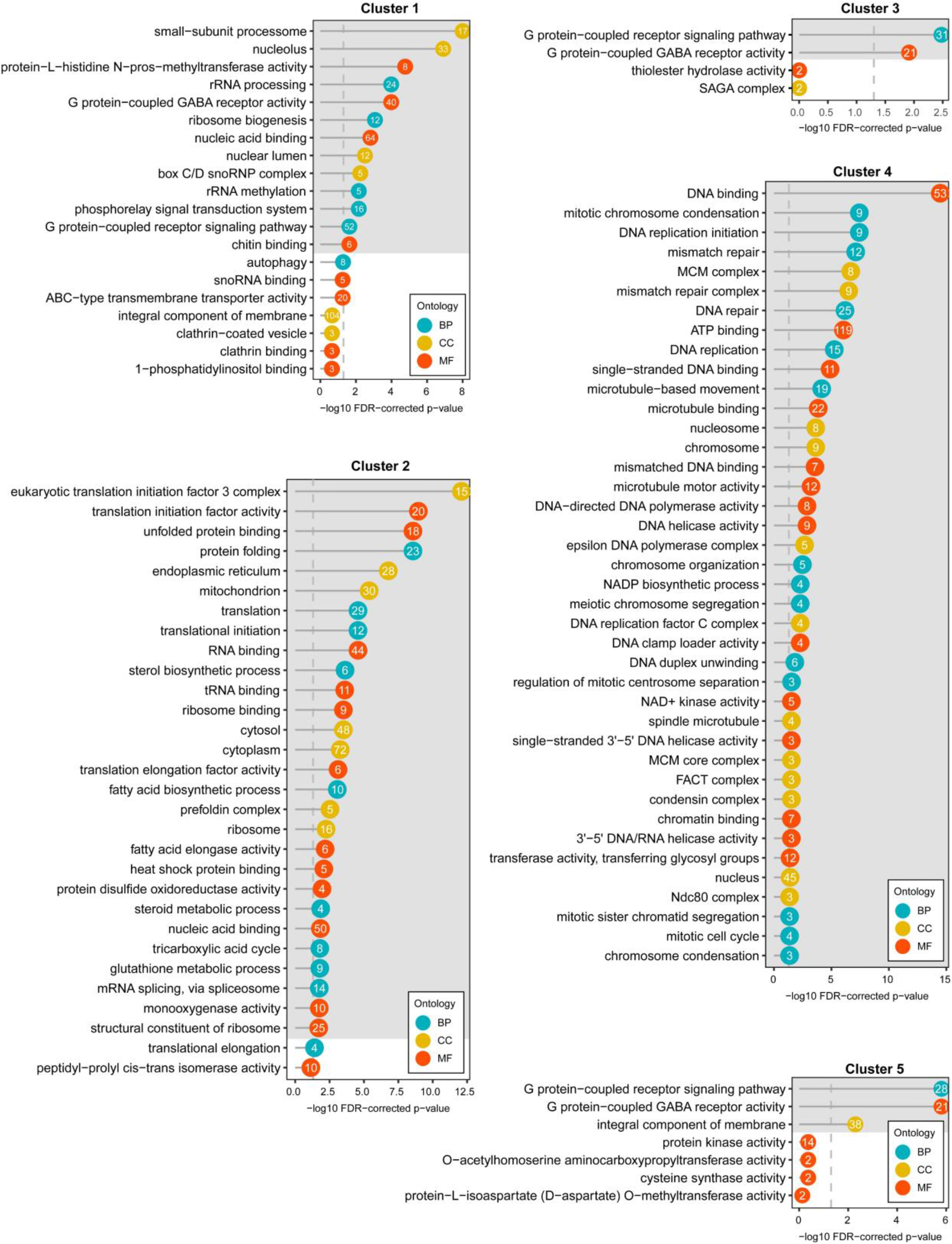
Enriched GO terms associated with each of the clusters. For each cluster of transcripts with similar expression profiles shown in Figure 2, GO terms are ranked according to the -log10 FDR-corrected p-value. A grey background corresponds to an FDR-corrected p-value cut-off of 0.05. The number in the dots indicates the number of contigs in the cluster that this term is associated with. The ontology of the respective term is shown in turquoise, (biological process, BP), yellow (cellular component, CC), or orange (molecular function, MF).

**Table S1:** Statistics of the rnaSPAdes and Trinity transcriptome assemblies.

|  | rnaSPAdes | Trinity |
| --- | --- | --- |
| Isoforms | 52,058 | 49,848 |
| Genes | 34,713 | - |
| ORFs | 47,527 | 49,848 |
| E90N50 | 2,698 | 1248 |
| Mean contig length | 1,805 | 871 |
| Median contig length | 1,192 | 612 |
| % GC | 44.91 | 45.85 |

**Figure S2:**
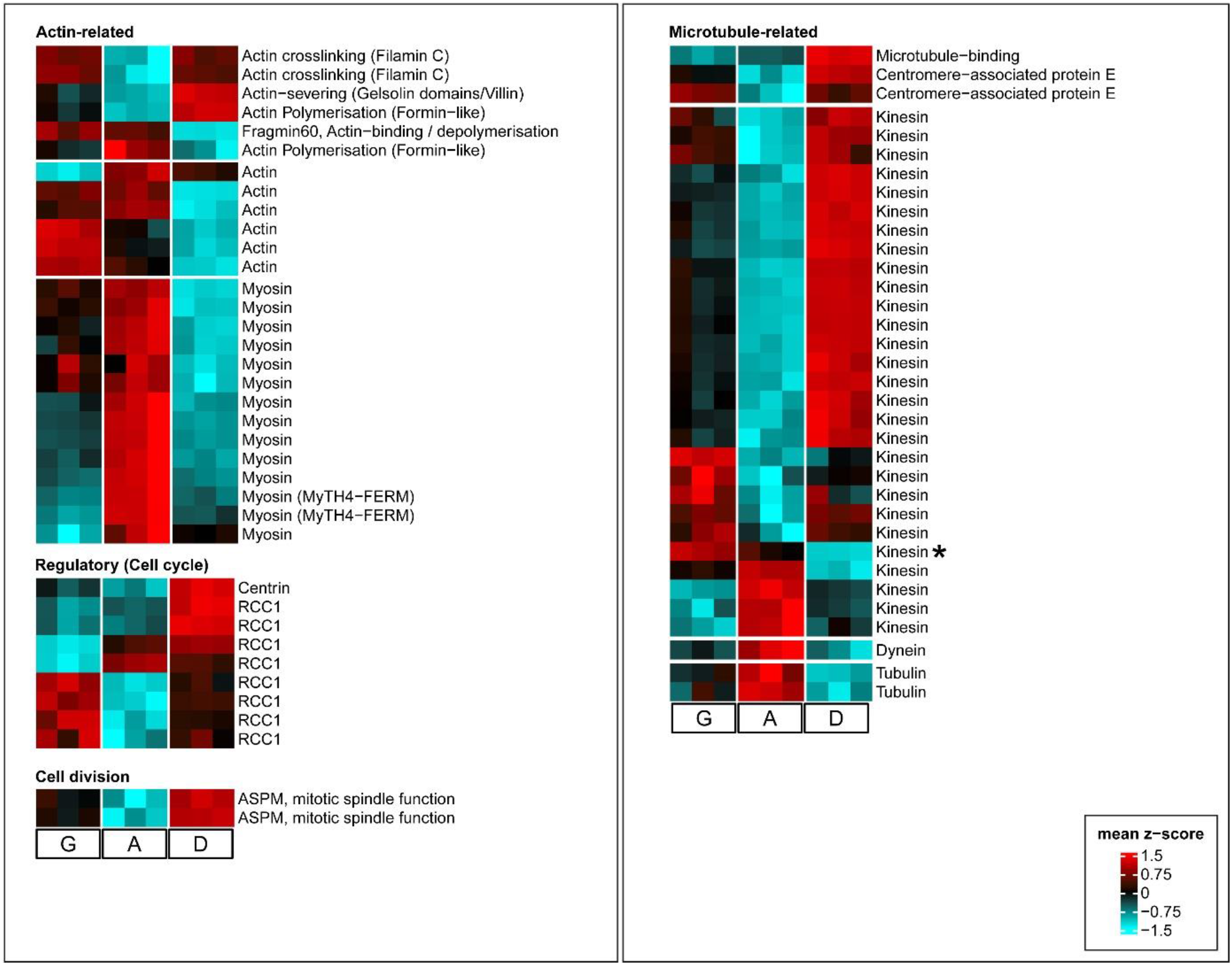
Heatmap of differentially expressed transcripts with a predicted cytoskeletal function (Functional eggNOG category “Z: Cytoskeleton”). G: Gliding, A: Attacking, D: Digesting. Each square represents one biological replicate. The asterisk marks the putative KIF17/OSM-3 homologue discussed in the main text.

**Figure S3:**
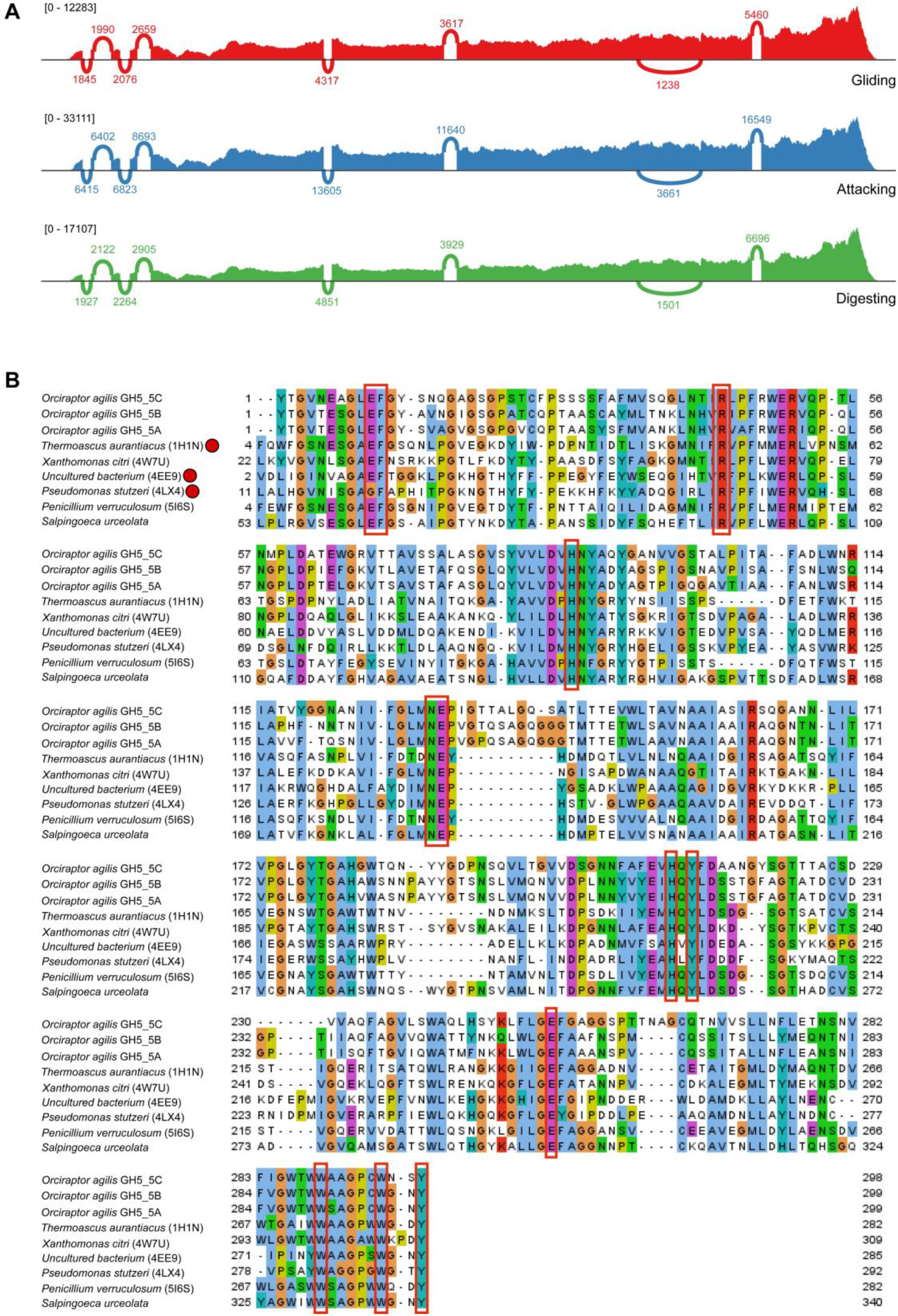
GH5_5 cellulases in *Orciraptor agilis*. **A**: Sashimi plot of the GH5_5A gene in the rnaSPAdes assembly. Shown is the coverage of reads along the sequence in each condition. Also indicated are the numbers of junction-spanning reads. Only junctions with more than 1000 reads are shown. **B**: Alignment of GH5_5 domains including the three GH5_5 sequences detected in *Orciraptor agilis*. Conserved residues that are part of the active site in known structures are marked with red boxes^15^. Functionally characterised glycoside hydrolases are marked with a red dot.

